# A computational probe into the behavioral and neural markers of atypical facial emotion processing in autism

**DOI:** 10.1101/2021.03.24.436640

**Authors:** Kohitij Kar

**Affiliations:** McGovern Institute for Brain Research and Department of Brain and Cognitive Sciences, Massachusetts Institute of Technology, Cambridge, MA, 01239, USA; Center for Brains, Minds, and Machines, Massachusetts Institute of Technology, Cambridge, MA, 01239, USA

**Keywords:** Autism, Amygdala, Inferior Temporal Cortex, Artificial Neural Networks, Facial emotion recognition

## Abstract

Despite ample behavioral evidence of atypical facial emotion processing in individuals with autism (IwA), the neural underpinnings of such behavioral heterogeneities remain unclear. Here, I have used brain-tissue mapped artificial neural network (ANN) models of primate vision to probe candidate neural and behavior markers of atypical facial emotion recognition in IwA at an image-by-image level. Interestingly, the ANNs’ image-level behavioral patterns better matched the neurotypical subjects’ behavior than those measured in IwA. This behavioral mismatch was most remarkable when the ANN behavior was decoded from units that correspond to the primate inferior temporal (IT) cortex. ANN-IT responses also explained a significant fraction of the image-level behavioral predictivity associated with neural activity in the human amygdala — strongly suggesting that the previously reported facial emotion intensity encodes in the human amygdala could be primarily driven by projections from the IT cortex. Furthermore, in silico experiments revealed how learning under noisy sensory representations could lead to atypical facial emotion processing that better matches the image-level behavior observed in IwA. In sum, these results identify primate IT activity as a candidate neural marker and demonstrate how ANN models of vision can be used to generate neural circuit-level hypotheses and guide future human and non-human primate studies in autism.

## Introduction

The ability to recognize others’ mood, emotion, and intent from facial expressions lie at the core of human interpersonal communication and social engagement. This relatively automatic, visuocognitive feature that neurotypically developed human adults take for granted shows significant differences in children and adults with autism ^1–4^. A mechanistic understanding of the underlying neural correlates of such behavioral mismatches is key to designing efficient cognitive therapies and other approaches to help individuals with autism.

There is a growing body of work on how facial identity is encoded in the primate brain, especially in the Fusiform Face Areas (FFA) in humans ^5,6^ and in the topographically specific “face patch” systems of the inferior temporal (IT) cortex of the rhesus macaques ^7–9^. Also, previous research has linked human amygdala neural responses with recognizing facial emotions ^10–12^. For instance, subjects who lack a functional amygdala often exhibit selective impairments in recognizing fearful faces ^13,14^. Wang et al.^15^ also demonstrated that the human amygdala parametrically encodes the intensity of specific facial emotions (e.g., fear, happiness) and their categorical ambiguity. A critical question, however, is whether the atypical facial emotion recognition broadly reported in individuals with autism (IwA) arises purely from differences in sensory representations (i.e., purely perceptual alterations^16,17^) or is due to a primary (but not mutually exclusive) variation in the development and function of specialized affect processing regions (e.g., atypical amygdala development leading to specific differences in encoding emotion). There are two main roadblocks toward answering this question. First, heterogeneity and idiosyncrasies are commonplace across behavioral reports in autism, including facial affect processing (for a formal meta-analysis of recognition of emotions in autism see: ^18,19^). The inability to parsimoniously explain such heterogeneous findings prevent us from designing more efficient follow-up experiments to probe such questions further. Second, in the absence of neurally mechanistic models of behavior, it remains challenging to infer neural mechanisms from behavioral results and generate testable neural circuit level predictions that can be validated or falsified using neurophysiological approaches. Therefore, we need brain-mapped computational models that can predict at an image-by-image level how primates represent facial emotions across different parts of their brain and how such representations are linked to their performance in facial emotion judgment tasks (like the one used in ^4^).

The differences in facial emotion judgments between neurotypical adults and individuals with autism are often interpreted with inferential models (e.g., psychometric functions) that base their predictions on high-level categorical descriptors of the stimuli (e.g., overall facial expression levels of “happiness”, “fear” and other primary emotions^20^). Such modeling efforts are likely to ignore an important source of variance produced by the image-level sensory representations of each stimuli being tested. To interpret this source of variance, it is necessary to develop models that are image computable. Recent progress in computer vision and computational neuroscience has led to the development of artificial neural network (ANN) models that can both perform human-like object recognition ^21,22^ as well as contain internal components that match human and macaque visual systems ^23,24^. Such image-computable ANNs can generate testable neural hypotheses ^25,26^ and help design experiments that leverage on the image-level variance to guide us beyond the standard parametric approaches.

In this study, I have used a family of brain-tissue mapped ANN models of primate vision to generate testable hypotheses and identify candidate neural and behavior markers of atypical facial emotion recognition in IwA. Specifically, I have compared the predictions of ANN models with behavior measured in neurotypical adults and people with autism ^4^, and facial emotion decodes from neural activity measured in the human amygdala ^15^. Furthermore, I performed in silico perturbation experiments to simulate and test autism-relevant hypotheses of underlying neural mechanisms. I observed that the ANNs could accurately predict the human facial emotion judgments at an image-by-image level. Interestingly, the models’ image-level behavioral patterns better matched the neurotypical human subjects’ behavior than those measured in individuals with autism. This behavioral mismatch was most remarkable when the model behavior was constructed from units that correspond to the primate IT cortex. Interestingly, I also observed this behavioral mismatch when comparing neural decodes from a distinct population of visually facilitated neurons in the human amygdala with *Control* and IwA behavior. However, ANN-IT activation patterns could fully account for the image-level behavioral predictivity of the human amygdala population responses that has been previously implicated in autism-related facial emotion processing differences ^12,15^. Furthermore, in silico experiments revealed that learning the emotion discrimination task with noisier ANN-IT representations (i.e., with higher response variability per unit) result in weaker synaptic connections between the model-IT and the downstream decision unit that improve the model’s match to the image-level behavioral patterns measured in the IwA. In sum, these results argue that noisier sensory representations in the primate inferior temporal cortex that drive a distinct population of neurons in the human amygdala is a key candidate mechanism of atypical facial emotion processing in individuals with autism — a testable neural hypothesis for future human and nonhuman primate studies.

## Results

As outlined above, I reasoned that the ability to predict the image-level differences in facial emotion judgments between individuals with autism (IwA) and neurotypical adults (*Controls*) allow us to 1) design more efficient experiments to study the atypical facial processing observed in IwA, 2) efficiently probe the underlying neural correlates. In this study, I first took a data-driven approach to discover such image-level differences in behavior across *Controls* and IwA in a facial emotion discrimination task ^4^. I then used brain-mapped computational models of primate vision to probe the underlying neural mechanisms that could drive such differences.

The behavioral and neural measurements analyzed in this study were performed by Wang et al. ^4,15^. During the task, participants were shown images of individual faces with specific levels of morphed emotions (for 1 sec) and asked to discriminate between two emotions, fear and happiness (Figure 1A; see Methods for details). The authors observed a reduced specificity in facial emotion judgment among individuals with autism (IwA) compared to neurotypical *Controls* (Figure 1B). Notably, the study controlled for low-level image confounds, and eye movement patterns across the two groups did not explain the reported behavioral differences. Therefore, the behavioral results significantly narrowed the space of neural hypotheses to sensory and affect-processing circuits.

**Figure 1.**
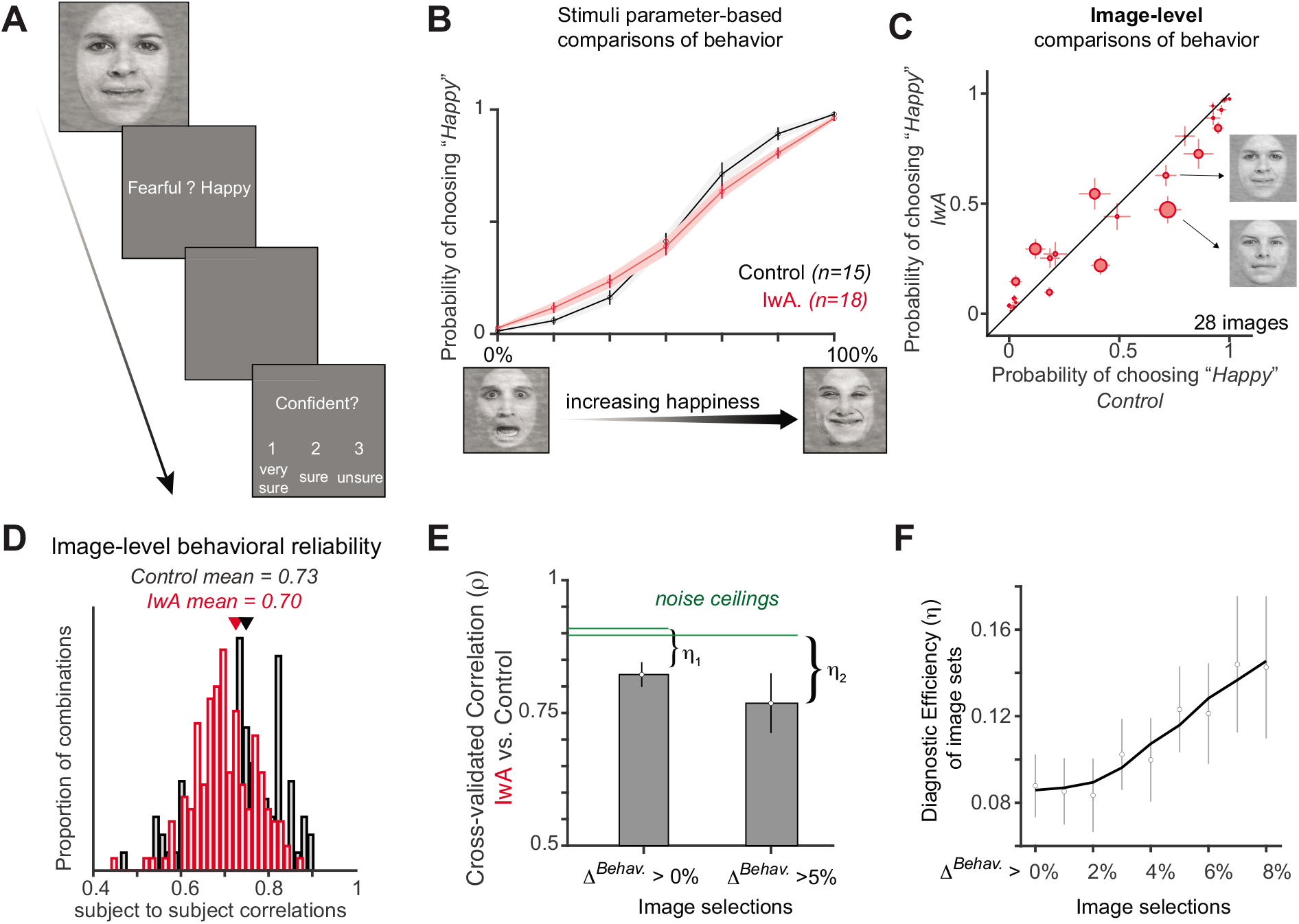
Behavioral task and image-level assessment of behavioral markers. **A.** Subjects, both neurotypical (*Control; n=15*) population and individuals with autism (IwA; n=18) viewed a face for 1 sec in their central ~12 deg, followed by a question asking them to identify the facial emotion (fearful or happy). After a blank screen of 500 ms, subjects were then asked to indicate their confidence in their decision (‘1’ for ‘very sure’, ‘2’ for ‘sure’ or ‘3’ for ‘unsure’). **B**. The psychometric curves show the proportion of trials judged as “happy” as a function of facial emotion morph levels (ranging from 0% happy (100% fearful; left) to 100% happy (0% fearful; right)). IwA (red curve), on average, showed lower specificity (slope of the psychometric curve) compared to the *Controls* (black curve). The shaded area and errorbars denotes SEM across participants. **C.** Image-level differences in behavior between *Controls* vs. IwA. Each red dot corresponds to an image. The size of the dot is scaled by the difference in behavior between the *Controls* and IwA. Errorbars denote SEM across subjects. Two example images are highlighted that show similar emotional (“happiness”) judgments by the *Controls* but drive significantly different behaviors in IwA — demonstrating the importance of investigating individual image-level differences. **D.** The estimated image-by-image happiness judgments were highly reliable as demonstrated by comparisons across individuals (estimated separately for each group). The mean reliability (average of the individual subject to subject correlations) was 0.73 and 0.70 for the *Controls* (black histogram) and IwA (red histogram), respectively. **E.** Correlation between image-by-image behavioral patterns measured in *Controls* vs. *IwA*, with two different selections of images (cross-validated image selections with held-out subjects). Noise ceilings were calculated based on measured behavioral (split-half) reliability across populations within each group (see Methods). The difference between the noise ceiling and the mean raw correlation is referred to as the diagnostic efficiency of the image-set (η) **F**. Diagnostic efficiency (*η*) as a function of image selection criteria. Errorbars denote bootstrap confidence intervals. Facial images shown in this figure are morphed and processed version of the original face images. These images have full re-use permission.

### Image-level differences can be leveraged to produce stronger behavioral markers of atypical facial emotion judgments in autism

Wang and Adolphs ^4^ primarily investigated the differences in behavior of IwA and *Controls*, across parametric variations of facial emotion levels (e.g., levels of happiness and fear). Here, I first examined whether the image-by-image behavioral patterns (irrespective of their facial identity or emotion levels), across the IwA and *Control* groups could be reliably estimated. Therefore, I computed the individual subject-to-subject correlations in image-level behavior (Figure 1D) which show that both of the groups exhibit highly reliable image-level behavior. The internal reliability (see Methods) for *Control* and IwA groups are 0.73 and 0.70, respectively. A visual inspection of the comparison of behavioral patterns across the two groups (Figure 1C) show that there are pairs of images (two such examples are shown in Figure 1C) for which the *Control* group exhibited very similar behavior, but the IwA made very different behavioral responses. This further confirms that diagnostic image-level variations in behavior could be further utilized to gain more insight into the mechanisms that drive the atypical facial emotion responses in IwA. Next, I quantified how stimuli selection based on high image-level differences can be leveraged to design more efficient behavioral experiments. To do this, I selected images based on the difference in behavior between the two groups (*Δ^Behav^*: using data from four randomly selected individual subjects from each group) and tested the resulting correlation between the two groups’ behavior (using the held-out subject population). This was repeated several times to get a mean measure of the cross-validated raw correlation (y-axis in Figure 1E). A noise-ceiling was measured for each image-set selection based on image-level internal reliability of the held-out test population (see Methods). The difference between the noise ceiling and the raw correlation is referred to as the diagnostic efficiency *η* of the image-set, which is a measure of how efficient the image-set is in discriminating between the IwA and *Control* behavior. Figure 1F shows how *η* varies across more and more efficient selection of image-sets (based on higher differences in image-level behavior with *Controls* and IwA). These results suggest that one reasonable goal of the field should be to find more efficient ways to predict which images will produce the highest *η* values. Focusing human behavioral testing on such images is likely going to yield stronger inferences and lead to a better understanding of the behavioral and neural markers driving the difference in behavior.

### ANN models of primate vision trained on varied objectives can perform facial emotion judgment tasks

To investigate how one can predict the image-level facial emotion judgments, I first tested how accurately current ANN models of primate vision can be trained to perform such tasks. One advantage of using these ANNs is that there are significant correspondences between their architectural components and the areas in the primate ventral visual cortex ^24,25,27^ (as shown in the schematic Figure 2A). Also, there is a significant match in the predicted behavioral patterns of such models with primate behavior (including face-related tasks) measured during multiple object recognition tasks^21,22^. Taken together, these models are great candidates for generating testable hypotheses regarding both neural and behavioral markers of specific visual tasks. I selected four different ANNs to test their behavioral predictions with respect to the facial emotional judgement task. These ANNs were pretrained to perform image classification (AlexNet^28^, CORnet-S^29^), face recognition (VGGFace^30^) and emotion recognition (Emotion-Net^31^). I observed that, a 10-fold cross validated partial least square regression model (see Methods for details) could be used to train each model to perform the task. The variation of the behavioral responses of the model with parametric changes in the level of happiness in the faces qualitatively matched the patterns observed in the human data (Figure 2B).

**Figure 2.**
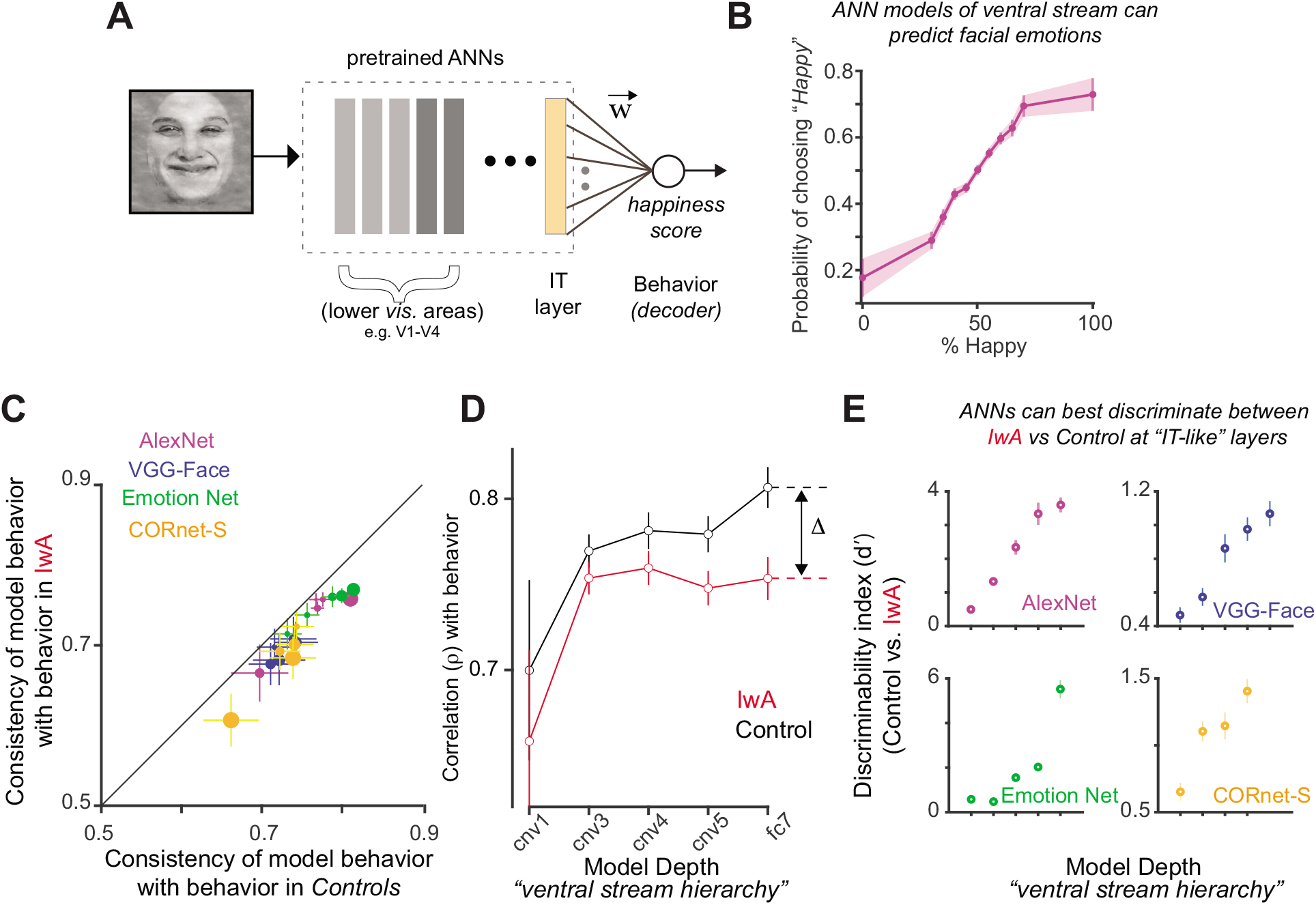
Testing ANN-models on facial emotion recognition tasks. **A.** ANN models of the primate ventral stream (typically comprising V1, V2, V4 and IT like layers) can be trained to predict human facial emotion judgments. This involves building a regression model, i.e., determining the weights 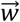 based on the model layer activations (as the predictor) to predict the image ground truth (“level of happiness”) on a set of training images, and then testing the predictions of this model on held-out images. **B.** An ANN model’s predicted psychometric curves (e.g., AlexNet, shown here) show the proportion of trials judged as “happy” as a function of facial emotion morph levels ranging from 0% happy (100% fearful; left) to 100% happy (0% fearful; right). This curve demonstrates that activations of ANN layers (layer ‘fc7’ that corresponds to the “model-IT” layer) can be successfully trained to predict facial emotions. **C.** Comparison of ANN’s image-level behavioral patterns with the behavior measured in *Controls* (x-axis) and IwA (y-axis). Four ANNs (with 5 models each generated from different layers of the ANNs are shown here in different colors. ANN predictions better match the behavior measured in the *Controls* compared to IwA. The correlation values (x and y axes) were corrected by the noise estimates per human population so that the differences are not due to differences in noise-levels in measurements across the IwA and *Control* subject pools. The dot size refers to the degree of discrepancy between ANN predictivity of *Controls* vs. IwA. **D**. A comparison of the ANN predictivity (results from AlexNet shown here) of behavior measured in IwA vs. *Controls* as function of model layers (convolutional (cnv) layers 1,3,4, and 5 and the fully connected layer 7, ‘fc7’ -- that approximately corresponds to the ventral stream cortical hierarchy). The difference between the ANN’s predictivity of behavior in IwA and *Controls* increases with depth and is referred to as Δ **. E.** Discriminability index (d’; ability to discriminate between image-level behavioral patterns measured in IwA vs. *Controls*; see Methods) as a function of model layers (all four tested models shown separately in individual panels). The difference in ANN predictivity between *Controls* and IwA was largest at the deeper (more IT-like) layers of the models instead of earlier (more V1, V2, and V4-like) layers. Errorbars denote bootstrap confidence intervals. Facial images shown in this figure are morphed and processed version of the original face images. These images have full re-use permission.

### ANN model predictions better match the behavioral patterns measured in neurotypical adults compared to individuals with autism

Next, I quantified how well the ANNs can predict the human image-level behavioral responses (across both *Controls* and IwA). Interestingly, ANN models significantly better predicted the image-level behavior measured in *Control* compared to the behavior measured in IwA (Figure 2C; 20 models tested; paired t-test; p<00001; t(19) = 10.99). To dissect which layer of the ANN best discriminated between the behavior of *Controls* and IwA, I compared individual models constructed from different layers of the same pretrained ANN architectures. This revealed two critical points. First, the correlation between model behavior and the *Control* group behavior increased as a function of model depth (black line; e.g. AlexNet shown in Figure 2D), which corresponds to the ventral visual hierarchy as reported in many studies^23,24^. Second, the difference in the model’s predictivity of behavior measured in Controls vs. IwA across layers is also highest at deeper layers, which corresponds to primate IT (comparison of the black and the red line for AlexNet shown in Figure 2D). This overall qualitative observation was consistent across all four tested models (Figure 2E). Given the high discriminability index (see Methods), established mappings between the layers and primate brain, as well as wide usage among researchers, I have used AlexNet for the subsequent analysis presented in this study. Therefore, these results suggest that population neural activity in primate IT could play a significant role in the atypical facial emotion processing in people with autism, and the image-level differences in sensory representations in IT might explain the difference in behavior observed across the images. However, such a role has been previously attributed to the human amygdala responses ^15^. Therefore, I next tested whether the human amygdala responses can predict the image-level behavior and how well this predictivity could be explained by the ANN-IT representations.

### Two distinct neural population coding schemes in the human amygdala

Wang et al.^15^ recorded bilaterally from implanted depth electrodes in the human amygdala (schematic shown in Figure 3A) from patients with pharmacologically intractable epilepsy. Subjects were presented each image for 1s (same as the task description above^4^) to discriminate between two emotions, fear and happiness. Similar to previous reports^15^, I observed two distinct population of neurons in the human amygdala. These two populations were marked by significant response suppression (visually suppressed (VS); 57 neurons; Figure 3B, right panel) and facilitation (visually facilitated (VF); 99 neurons; Figure 3B, left panel) respectively, after the onset of the facial image stimulus. I first tested how well the population-level activity (250-1500 ms post image onset) of three specific subsamples of the amygdala neurons (VS only, VF only and VS + VS neurons) predicted the behavioral patterns measured in human subjects. I observed that each of these populations of VF, VS, and mixed (equal number of VS and VF neurons) could significantly (p<0.0001; permutation test for significance of correlation) predict the image-level facial emotion judgments measured in *Controls*. Figure 3C shows how these three populations predict the image-level behavior measured in *Controls* as a function of the number of neurons sampled to build the neural population decoders. Given that all of these groups exhibit an increase in behavioral predictivity with the number of neurons, it is difficult to reject any of these decoding models (with the current neural dataset). Therefore, in the following analyses I have examined the VF and VS units separately. Next, I estimated how well the VS and VF population predicted the behavioral patterns measured in the *Control* and IwA respectively. Interestingly, I observed that similar to the ANN-IT behavior, neural decodes out of the VF neurons in the human amygdala better match the *Control* group behavior compared to the ones measured in IwA (Figure 3C; Δ*^VF^* is significantly greater than 0; permutation test of correlation: p<0.05). However, the VS neurons did not show this trend (Figure 3D; Δ*^VS^* is not significantly different from 0; permutation test of correlation; p>0.05). Figure 3E shows how VF (and not VS) neurons become more discriminatory of the IwA vs. *Control* behavior (i.e., Δ*^VF^* increases) as we choose image-sets with higher diagnostic efficiencies (η). Consistent with prior work, these results provide evidence that neural responses in the human amygdala are implicated in atypical facial processing in people with autism. However, the results presented here also critically identify the VF neurons as a stronger candidate neural marker of the differences in facial emotion processing observed in IwA.

**Figure 3.**
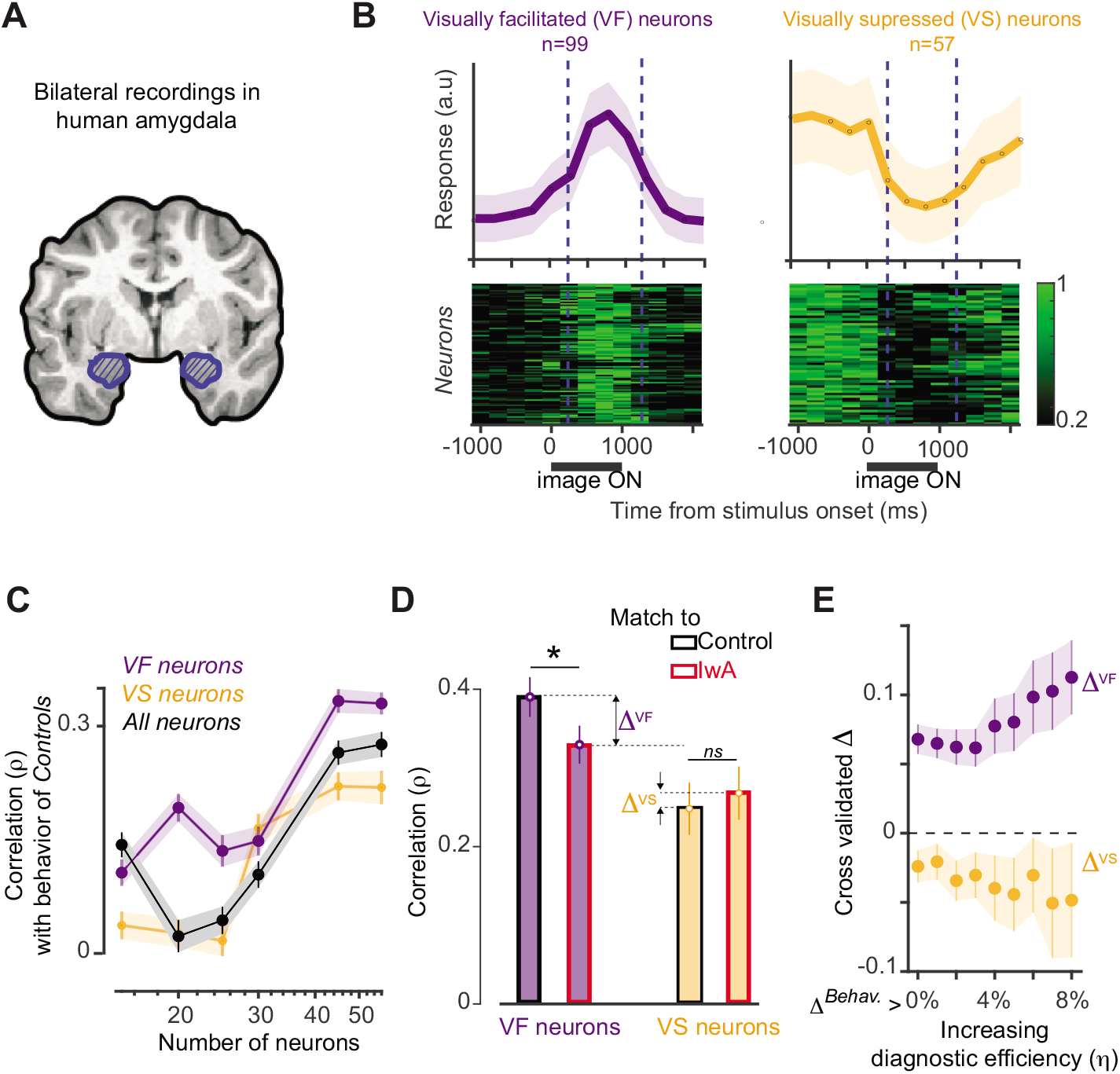
Facial emotion representation in the population neural activity of human amygdala. **A.** Schematic of bilateral amygdala (blue patch) recordings performed by Wang et al. **B.** Two distinct population of neurons observed in the human amygdala. The visually facilitated (VF; shown in purple) neurons (n=99) increased their responses after the onset of the face stimuli (top left panel: averaged normalized spike rate across time; 250 ms time bins). The bottom left panel shows the normalized firing rate across time for each VF neuron. The visually suppressed (VS; shown in yellow) neurons (n=57) decreased their responses after the onset of the face stimuli (top right panel: averaged normalized spike rate across time; 250 ms time bins). The bottom right panel shows the normalized firing rates across time for each VS neuron. Errorbars denote SEM across neurons. **C.** An estimate (correlation) of how three subsamples of neural populations, VS (yellow), VF (purple) and VS+VF (‘All’, black) predict the image-level behavior measured in *Controls* as a function of the number of neurons sampled to build the neural decoders. Errorbars denote bootstrapped CI. **D.** Comparison of how well the VS (yellow bars) and VF (purple bars) neurons predict the behavior measured in *Controls* vs. IwA. The red and black edges denote the predictivity of IwA and *Controls* respectively. Δ*^VF^* and Δ*^VS^* are the differences in the human amygdala (neural decode) predictivity of facial emotion judgments measured in *Controls* and IwA from the VF and VS neurons respectively. Errorbars denote bootstrap CI. **E.** Δ*^VF^* and Δ*^VS^* as function of image selection (which is proportional to the diagnostic efficiency η estimated per image-set). The cross validation was done at the level of subjects for each image selection. Errorbars denote bootstrap CI.

### ANN-IT features can explain a significant fraction of the image-level behavioral predictivity of the human amygdala population

Given the significant predictivity of facial emotion judgments observed in the ANN IT layers and the presence of strong anatomical connections between primate IT and amygdala^32^, I further asked how much of the image-level predictivity estimated from the amygdala activity is likely driven by input projections from the IT cortex. To test this, I first asked (with a linear regression analyses; see Methods) how well the image-by-image behavioral predictions from the ANN-IT models (AlexNet-fc7 tested here) can explain the image-by-image neural decoding patterns estimated from the amygdala neurons (separately for VS and VF neurons). The residue of this analyses (see Methods) contained the variance in the amygdala decodes that was not explained by the predictions of the ANN-IT models. Therefore, the amount of variance in the measured behavioral patterns explained by this residue provides an estimate of how much of the behavior is purely driven by the amygdala responses independent of the image-driven sensory representations. Assuming a feedforward hierarchical circuit whereby the IT cortex drives the human amygdala and not the other way around, a lower percentage of explained variance (%EV) obtained after such an analysis should indicate that the source of the signal in amygdala is at least partially coming from the IT cortex. Interestingly, this analysis revealed that the behavioral predictivity (%EV) of the human amygdala is significantly reduced once I regressed out the variance that is driven by the ANN-IT responses. For instance, when considering all images (i.e., very low diagnostic efficiency of the imageset), I observed that VS and VF neurons could explain approximately 17.24% and17.39% (a lower bound of the %EV since neural noise has not been accounted for) of the behavioral variance (Figure 4A, B; left panel). However, once the ANN-IT driven variance was regressed out these values significantly dropped to 0.06% and 0.2% respectively (Figure 4A, B; right panel). Overall, VF neural residuals (after regressing out ANN-IT predictions) explained significantly less variance at all tested η levels. VS neural residuals explained significantly less variance only at lower η levels (Δ*^Behav^* < 2.5%). Given that VS neurons showed a drop in %EV for higher η levels, it is not surprising that I did not observe any differences with the residual predictivity at those levels. Interestingly, there was no significant change in %EV across the image selections when VS activity was regressed out of VF activity (and vice versa; Figure 4A, B; middle panel), providing further evidence that they largely support a complimentary coding scheme for facial emotions within the amygdala. In sum, these results suggest that input projections from the IT cortex into the amygdala ^32^ might be the primary career of the facial emotion related signals. Furthermore, the results also suggest a likely difference in how VS and VF neurons are affected in IwA – with VF neurons being more diagnostic of the atypical behavior observed in IwA.

**Figure 4.**
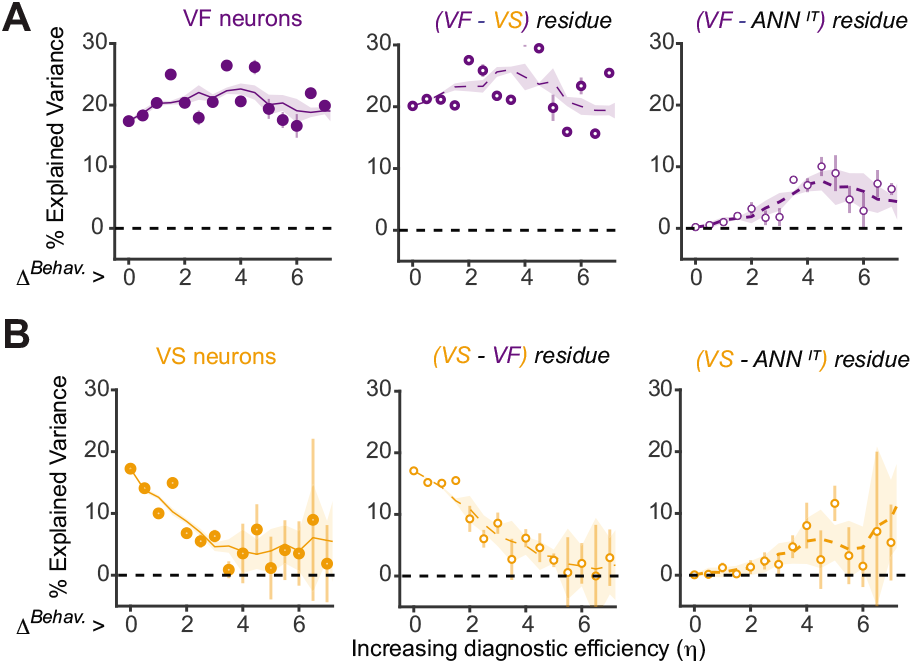
Amount of behavioral variance (measured in *Controls*) explained by different neural markers. **A.** Left panel: Percentage of behavioral variance explained by the human amygdala (VF) neural activity as a function of the overall differences in image-level behavior between IwA and *Controls*. As demonstrated in Figure 1F the x-axis is proportional to the diagnostic efficiency (η). Middle panel: Percentage of variance explained by the residual (VS-based predictions regressed out of the predictions from VF-based neural decodes). There was no significant change in %EV across the image selections when VS was regressed out, suggesting a complimentary coding scheme. Right panel: Percentage of behavioral variance explained by the residual (ANN-IT predictions regressed out of the predictions from VF-based neural decodes). There was a significant difference (reduction in %EV) between the two cases for all levels of tested η. **B.** Left panel: Percentage of behavioral variance explained by the human amygdala (VS) neural activity as a function of the overall differences in image-level behavior between IwA and *Controls*. Middle panel: Percentage of variance explained by the residual (VF-based predictions regressed out of the predictions from VS-based neural decodes). There was no significant change in %EV across the image selections when VF was regressed out, suggesting a complimentary coding scheme. Right panel: Percentage of variance explained by the residual (ANN-IT predictions regressed out of the predictions from VS-based neural decodes). There was a significant difference (reduction in %EV) between the two cases while Δ*^Behav^* was less than 2. All %EV values were estimated in a cross validated way, wherein the image selections and the final estimates were done based on different groups of subjects. Errorbars denote bootstrapped CI.

### In silico perturbations with additional noise in ANN-IT layers improves the model’s match with the behavior of individuals with autism

To further probe how IT representations might be different in IwA compared to *Controls* (Figure 5A), I compared ANNs independently trained to predict the behavior of *Controls* and IwA. I directly compared the learned weights, that is the synaptic strengths between the model-IT layer and the behavioral output node in the two cases. I observed that are model trained on the behavior measured in IwA yielded weaker synaptic strengths for both excitatory (positively weighted) and inhibitory (negatively weighted) connections (Figure 5B), compared to models trained to reproduce the behavior measured in *Controls*. I further explored how this modest difference in the models could be simulated such that an ANN trained on ground truth labels of human facial emotions could be transformed into behaving more like what we observe in IwA. Based on previous studies ^33,34^, I hypothesized that increased noise (scaled according to overall responsiveness of the model units) in the sensory representations during learning could potentially yield weaker synaptic strengths between the model-IT layer and the trained behavioral output node. Of note, although a noisy representation likely yields a reduced specificity in behavioral performance, an addition of specific amounts of noise does not necessarily guarantee a stronger or weaker correlation with the image-level behavioral patterns observed in IwA. Therefore, such in silico perturbations could produce three primary outcomes. First, adding noise might produce no effects in the model’s behavioral match with the behavior of IwA (Figure 5C, top panel, *H*_0_). Second, the added noise might weaken the correlation achieved by a noiseless model (Figure 5C, middle panel, *H*_1_). Third, and consistent with an Autism Spectrum Disorder ASD)-relevant mechanism, addition of noise could improve the correlation with the image-level behavior measured in IwA (Figure 5C, bottom panel, *H*_2_). I observed that at specific levels of added noise (Figure 5D; dashed black line) during the model training (transfer learning), the model’s behavioral match with IwA significantly improved (assessed by permutation test of correlation) beyond the levels noted with a noise-free model (Figure 5D). In addition, this increase in the predictivity of IwA behavior with addition of noise is significantly higher than that observed when compared to the model’s predictivity of the behavior measured in the *Controls* (as shown in Figure 5E). Within the dashed black lines (Figure 5E), noise added to each model unit were drawn from a normal distribution with zero mean and standard deviation equal to 2 to 5 times the width of the response distribution of that unit across all tested images. Taken together, this strongly suggests that additional noise in sensory representations is a very likely candidate mechanism implicated in atypical facial emotion processing in adult with autism.

**Figure 5.**
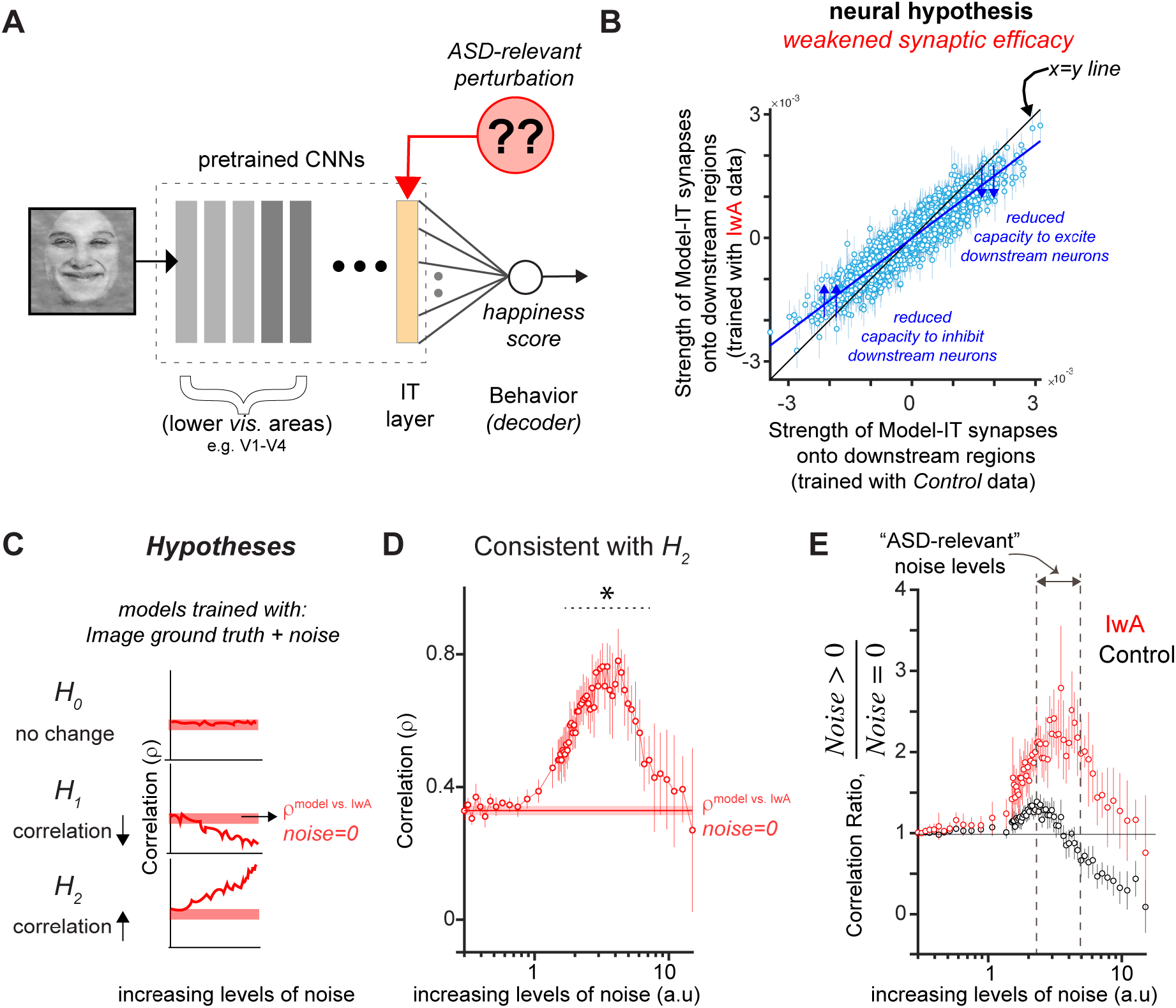
In silico experiments on ANNs to probe neural mechanisms underlying atypical facial emotion judgments in individuals with autism. **A.** What changes can one induce in the model-IT layer to simulate the behavioral patterns measured in IwA? **B.** Comparison of synaptic strengths (weights) between ANN-IT and the behavioral node when models are independently trained with the behavior measured in IwA vs. *Controls*. ANN fits to behavior of IwA yielded weaker synaptic strengths for both excitatory (positively weighted) and inhibitory (negatively weighted) connections. Each blue dot refers to the weights in the connection between an individual model unit in the IT-layer and the decision (“level of happiness”) node. **C.** Hypotheses and corresponding predictions *H*_0_: Addition of noise could lead to no differences in how it affects the model’s match to behavior measured in IwA. *H*_1_: Addition of noise could reduce the models’ match to behavior measured in IwA compared to the noise-free model. *H*_2_: Addition of noise could improve the models’ match to the behavior measured in IwA compared to the noise-free model. *H*_2_ supports the “high IT variability in autism” hypotheses. **D**. Correlation of ANN behavior with IwA as a function of levels of added noise. The results show that at specific noise regimes ANNs are significantly more predictive of the behavior measured in IwA compared to the noiseless model. Errorbars denote bootstrapped CI. **E.** Ratio of ANN behavioral predictivity of noisy vs. noise-free ANNs. At specific levels of noise, referred to as the Autism Spectrum Disorder (ASD)-relevant noise levels, the ANNs trained with noise show much higher predictivity for behavior measured in IwA while suffering a reduction in predictivity of the *Controls*. Errorbars denote bootstrapped CI. Facial images shown in this figure are morphed and processed version of the original face images. These images have full re-use permission.

## Discussion

The overall goal of this study was to identify candidate neural and behavioral markers of atypical facial emotion judgments observed in individuals with autism. Based on discovering reliable image-by-image differences between the behavior of *Controls* and IwA that could not be explained by categorical ambiguity in the stimuli, I reasoned that such image-level variance could be leveraged to probe the neural mechanisms of behavioral differences observed in IwA. Therefore, I used image-computable, brain-tissue mapped artificial neural network models of primate vision to further probe the issue. By using computational models (that have established brain tissue correlates) to explain experimental data, I hereby demonstrate how such an approach could be used to probe the neural mechanisms that underlie the differences in facial emotion processing observed in individuals with autism. Below, I discuss the findings with their relevance to future experiments and candidate mechanisms implicated in atypical facial emotion recognition in IwA.

### ANN based predictions can be used to efficiently screen images and provide neural hypotheses for more powerful experiments

A family of ANN models can currently predict a significant amount of variance measured in various object recognition related behaviors and neural circuits ^35^. Given that the results presented here demonstrate the ability of such ANNs to discriminate between the behavior measured in *Controls* and IwA, we can further leverage the ANNs to screen facial image stimuli and select images where the predicted behavioral differences are maximum. Further, such models can be reverse engineered^25,36^ to synthesize images that could achieve maximum differences to optimize behavioral testing and diagnosis. Such deep image synthesis methods could also modify the facial images such that the differences in the observed behavior between the *Controls* and IwA are minimized. Although clearly at an early stage, such methods have a significant potential to improve future cognitive therapies. Unlike many machine learning approaches that are not closely tied to the computation and architecture of the primate brain, the ANNs used in this study have established homologies with the primate brain and behavior ^35^. As demonstrated in this study, these links allow us to relate the ANN predictions to distinct brain areas directly. Specifically, the ANN results presented here suggest that population activity patterns in areas like the human and macaque inferior temporal cortex are vital candidates for neural markers of atypical facial processing in autism. The modeling results provide further insights into the most affected aspects of the population responses, implicating noisier sensory representations (see below) as a source of the differences in sensory representation, learning and subsequent decision making. Besides the specific hypotheses generated in this study, it is essential to note that ANN models of primate vision are an active area of research, and we are witnessing the gradual emergence of better brain-matched models ^29,37–39^. Therefore, this study establishes a critical link between atypical face processing in autism and how to leverage ANNs to study this.

### Modeling results imply the need for more fine grain neural measurements in the primate IT cortex and amygdala

The ANN-based computational analyses in this study provide specific neural hypotheses that can be tested using macaque electrophysiology and human fMRI experiments. First, I observed that the ANN-IT layers could best discriminate between the behavior of *Controls* vs. IwA. Therefore, such signals are likely also measurable in the primate IT cortex and are key candidates for neural markers of atypical facial emotion processing in autism. Given that most ANN models are feedforward-only or have minimal dynamics, it will be critical to test how the different temporal components of IT population responses carry the facial emotion signal. Similar to predictions of ANN-IT layers, I observed that population activity in the human amygdala also better matches behavior measured in the *Controls* than IwA. There can be multiple reasons for the observed differences in behavioral predictivity. First, it is possible that due to the atypical development of the human amygdala in IwA, the behavior they exhibit does not match well with the neural decodes out of the neurotypical amygdala. Second, the lack of predictivity might be carried forward from responses in the IT cortex -- as predicted by the ANNs. The current study attempted to disambiguate between these two factors. I asked how well ANN-IT predictions can account for the amygdala activity’s behavioral patterns. Indeed, the image-level predictivity of facial emotion judgments observed in the human amygdala’s population activity (both VF and VS neurons) was significantly explained away by the ANN-IT features (Figure 4A, B; left panel). This result is consistent with the hypothesis that the higher-level visual cortices (like IT) primarily drive the facial affect signal observed in the human amygdala. Simultaneous neural recordings in IT and amygdala or finer grain causal perturbation experiments need to be conducted to test this hypothesis more directly. Notably, the behavioral mismatch (neural decodes vs. *Control/IwA* behavior) was specific to the decodes constructed from the VF neurons (and not VS neurons). Therefore, future experimental investigations should dissect the role of IT cortex and how it functionally influences the VF and VS neurons, which are likely part of a complimentary coding scheme. Furthermore, it will be essential to examine how the IT cortical activity is driven by feedback projections from the amygdala, given that evidence for the importance of such connections from ventrolateral PFC has been demonstrated for object recognition^40^.

### High variability in sensory representation can lead to weaker efferent synaptic strengths during learning and development

In a psychophysical discrimination task, the typical consequence of having a noisy detector is a reduction in the sensitivity of performance, which manifests as a reduced estimated slope of the psychometric function. This is consistent with what Wang and Adolphs ^4^ had observed. Given that the idea of higher sensory variability in autism is also consistent with previous findings^34^, I considered this as a potential neural mechanism that could explain the image-level differences I have observed in the facial emotion discrimination behavior in IwA. Therefore, I tested the “increased sensory noise hypothesis” to test whether such a perturbation could simulate the weaker efferent synaptic connections from IT-like layers as revealed by the ANN based analyses (Figure 5B). Indeed, addition of noise during learning made the ANN behavior more matched with that observed in IwA. First, this could suggest that perhaps the behavior measured in IwA results from additional noise in the sensory representations that affects the subjects’ behavior during the task. However, this could also be the result of executing an inference engine (in the brain) that learned its representations under high sensory noise during development (as a child). An estimate of noise levels (sensory cortical signal variability) in children with autism and a quantitative probe into how that could potentially interact with learning new tasks is essential to test this hypothesis. As demonstrated in this study, the ANN models provide a very efficient framework to generate more diagnostic image-sets for these future studies given that we can simulate any level (and type) of noise under different learning regimes and make predictions on effect sizes. Such model-driven hypotheses are likely to play a vital role in guiding future experimental efforts and inferences.

### High variability in sensory representation can qualitatively explain other ASD-specific behavioral reports

Addition of noise during the transfer learning procedure of the ANN models made the model’s behavioral output more consistent with the behavior measured in IwA (Figure 5D). Such a mechanism can indeed qualitatively explain other previous behavioral observations made in individuals with autism. For example, Behrmann et al.^41^ observed that reaction times measured during object discrimination tasks, in adults with autism were significantly higher than the *Control* subjects. This difference was especially high during more fine-grained discrimination tasks. Such a behavioral phenomenon can be explained by an increase in sensory noise in IwA that leads to longer time requirements during integration of information ^42^, and weaker performances on finer discrimination tasks. The ANN based approach demonstrated in this study, however, provides guidance beyond the qualitative predictions of overall effect types. Specific image-level predictions provided by ANNs will help researchers to design more diagnostic behavioral experiments and make measurements that can efficiently discriminate among competing models of brain mechanisms.

### Potential underlying mechanisms behind increased neural variability

An imbalance in the ratio of the excitatory and inhibitory processes in cortical circuits has been proposed as an underlying mechanism for various atypical behaviors observed in autism^43^. I speculate that such an E/I imbalance could arise due to lower inhibition in the cortical networks. This could lead to larger neural variability and a subsequent noisier, less efficient sensory processing. Therefore, the results observed in the in-silico experiments are not biologically implausible. In fact, genetic mutations that impact the generation and function of interneurons have been previously linked with autism^44,45^. Therefore, cell-type specific causal perturbation approaches are necessary to test whether a decreased inhibition in the visuocortical pathway (especially in the primate IT cortex) leads to noisier sensory representations and can reproduce the specific image-level differences in facial emotion processing reported in this study. The image-level behavioral measurements and ANN predictions reported here will enable such stronger forms of hypothesis testing during the interpretation of such experimental results.

## Methods and Materials

### Human Behavior

In this study, I have re-analyzed behavioral data that was previously collected and used in a study by Wang and Adolphs^4^. The raw behavioral dataset was kindly shared via personal communication.

#### Participants

In the original study (for further details see^4^), eighteen high-functioning participants with ASD (15 male) were recruited. All ASD participants met DSM-V/ICD-10 diagnostic criteria for autism spectrum disorder (ASD) and met the cutoff scores for ASD on the Autism Diagnostic Observation Schedule-2 (ADOS-2) revised scoring system for Module 4, and the Autism Diagnostic Interview-Revised (ADI-R) or Social Communication Questionnaire (SCQ) when an informant was available. The ASD group had a full-scale IQ (FSIQ) of 105±13.3 (from the Wechsler Abbreviated Scale of Intelligence-2), a mean age of 30.8±7.40 years, a mean Autism Spectrum Quotient (AQ) of 29.3±8.28, a mean SRS-2 Adult Self Report (SRS-A-SR) of 84.6±21.5, and a mean Benton score of 46.1 ±3.89 (Benton scores 41–54 were in the normal range). ADOS item scores were not available for two participants, so we were unable to utilize the revised scoring system. But these individuals ‘original ADOS algorithm scores all met the cutoff scores for ASD.

Fifteen neurologically and psychiatrically healthy participants with no family history of ASD (11 male) were recruited as *Controls. Controls* had a comparable FSIQ of 107±8.69 (two-tailed t-test, P=0.74) and a comparable mean age of 35.1±11.4 years (P=0.20), but a lower AQ (17.7±4.29, P=4.62×10^-5^ and SRS-A-SR (51.0±30.3, P=0.0039) as expected. Participants gave written informed consent, and all original experiments were approved by the Caltech Institutional Review Board. All participants had normal or corrected-to-normal visual acuity. No enrolled participants were excluded for any reasons.

#### Facial emotion judgment task

During the task, Wang and Adolphs^4^ asked participants to discriminate between two emotions, fear and happiness. The image-set includes faces of four individuals (2 female) each posing fear and happiness expressions from the STOIC database (Roy et al. 2007), which are expressing highly recognizable emotions. To generate the morphed expression continua for the experiments, the authors interpolated pixel value and location between fearful exemplar faces and happy exemplar faces using a piece-wise cubic-spline transformation over a Delaunay tessellation of manually selected control points. They created 5 levels of fear-happy morphs, ranging from 30% fear/70% happy to 70% fear/30% happy in steps of 10% (Figure 1B). Low-level image properties were equalized using the SHINE toolbox ^46^. In each trial, a face was presented for 1 second followed by a question prompt asking participants to make the best guess of the facial emotion (Figure 1A). After stimulus offset, participants had 2 seconds to respond, otherwise the trial was aborted and discarded. Participants were instructed to respond as quickly as possible, but only after stimulus offset. No feedback message was displayed, and the order of faces was completely randomized for each participant. Images were presented approximately in the central 1° of visual angle. A subset of the participants (11 participants with autism and 11 *Controls*) also performed confidence ratings after emotion judgment and a 500 ms blank screen, participants were asked to indicate their confidence by pushing the button ‘1 ‘for ‘very sure’, ‘2 ‘for ‘sure ‘or ‘3 ‘for ‘unsure’. This question also had 2 seconds to respond. All images used in this study has free re-use permission as set here^15^.

#### Estimating image-level behavioral reliability

To estimate the image-level behavioral reliability (Figure 1D), I first estimated the probability of choosing “Happy” per image in each subject (15 Controls, 18 IwA) -- referred to as the 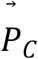 and the 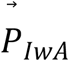 vectors. Then, for each possible combination of selecting 2 subjects from the subject pools, I estimated the subject-to-subject Kendall rank correlation coefficient. This was done separately for the Controls and IwA, leading to the red and black histograms in Figure1D respectively. These correlations scores are not corrected by the individual subjects’ internal reliability (across trials). Therefore, they represent the lower bound of the inter subject correlations.

#### Estimating noise ceilings for IwA vs. *Control* correlations

I define the noise ceiling of a correlation as the highest possible value of correlation expected given the noise measured independently in the two variables that are being tested. To estimate this, first I individually estimate the split half reliability of the 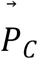 and the 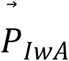 vectors. Each split is constructed with a random sampling of half of the subjects and taking the average across them and doing same for the other half of the subjects. For each iteration, such splits were made, and the correlation between the resulting vectors was computed. This correlation score was corrected by the Spearman-Brown correction procedure to account for the halving of subject numbers. I then computed the average across 100 such iterations, referred to as 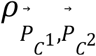 and 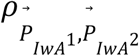 for the *Controls* and IwA respectively. The noise ceiling was then estimated as,

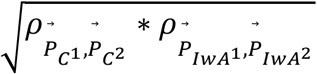

Intuitively, if both groups provided noiseless data, then these reliabilities should be each at 1, and therefore the noise ceiling shall also be set at 1. Noisy data will lead to <1 values for the individual 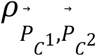 and 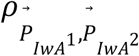 reliabilities, and hence the noise ceiling shall also be <1. Of note, each selection of image with result in a different 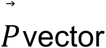 and therefore will result in a slightly different noise ceiling estimate, as demonstrated in Figure 1E (two green lines).

#### Estimating cross-validated diagnostic efficiency (*η*) of image-sets

Diagnostic Efficiency (*η*; shown in Figure1E, and 1F) of an image-set is defined as the cross-validated estimate of the difference between the noise ceiling and the raw correlation between the 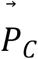 and the 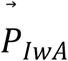 vectors. The cross validation is achieved by the choosing the images based on a specific subset of subjects and then measuring the noise ceiling and the raw correlation on a different held-out set of subjects. For efficient collection of human subject data that could optimally discriminate between the behavior measured in *Controls* and IwA, one must aspire for the highest *η* values for image-sets.

### Depth recording in human amygdala

In this study I have re-analyzed the neural data that was previously collected and used in a study by Wang et al.^15^. The raw neural dataset was kindly shared via personal communication. Wang and colleagues recorded bilaterally from implanted depth electrodes in the amygdala from patients with pharmacologically intractable epilepsy. Target locations in the amygdala were verified using post-implantation structural MRIs. At each site, they recorded from eight 40 *μ*m microwires inserted into a clinical electrode. Bipolar wide-band recordings (0.1–9 kHz), using one of the eight microwires as reference, were sampled at 32 kHz and stored continuously for off-line analysis with a Neuralynx system (Digital Cheetah; Neuralynx, Inc.). The raw signal was filtered with a zero-phase lag 300-3000 Hz bandpass filter and spikes were sorted using a semiautomatic template matching algorithm. Units were carefully isolated and spike sorting quality were assessed quantitatively. Subjects were presented each image for 1s (similar to the task description above) to discriminate between two emotions, fear and happiness.

#### Selection of neurons for analyses

In the original study, only units with an average firing rate of at least 0.2 Hz (entire task) were considered. Only single units were considered. In addition to that, in this study I have further restricted the neural dataset to neurons that have a significant visual response (both increase and decrease). To estimate that I compared the neural firing rates (per image) averaged across two specific time bins, [-1000 0] and [250 1250], where 0 is the onset of the image. If the paired Wilcoxon Signed Rank test between these two firing rate vectors were significant, the site was considered for further analyses. Thus, I considered 156 total neurons: 99 visually facilitated (VF) neurons and 57visually suppressed (VS) neurons.

#### Decoding facial emotion judgment from neural population activity

To decode facial emotion judgments from the neural responses per image, I used a linear model that linked the neural responses to the levels of happiness (ground truth from image generation). Building the model, essentially involves solving a regression problem estimating the weights 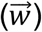 per neuron and a *bias* term. I used a partial least squares (MATLAB command: *plsregress*) regression procedure, using 15 retained components. I also used 10-fold cross validation. For each fold, the model was trained (i.e., 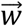 and *bias* were estimated) using the data from the other 9 folds (training data), and predictions were generated for the held-out fold (test images). This was repeated for each of the folds and the entire procedure was repeated 100 times. The predictions of the trained neural model on the held-out test images were used for future correlation analyses. Given the training scheme, every image was assigned as the test-image once per iteration.

### ANN models of primate vision

The term “model” in this study always refer to a specific modification of a pre-trained ANN. For instance, I have used an Image-Net pretrained deep neural network, AlexNet to build multiple models. Each model was constructed by deleting all layers succeeding a given layer. For instance, the ‘*cnv5*’ model was built by removing all layers of AlexNet that followed the output of its fifth convolutional layer. The feature activations from the fifth convolutional layer output were then trained with the linear regression procedure (similar to the neural decodes).

#### Estimating model facial emotion judgment behavior

To decode facial emotion judgments from the model responses per image, I used the same linear modeling approach as the neural data (see above), that linked the model feature activations to the level of happiness (ground truth from image generation). The model features, per layer, were extracted using the MATLAB command *activations* for AlexNet^28^, VGGFace^30^ and EmotionNet^31^ in MATLAB-R 2020b. For the CORnet-S^29^ model, I used the code from: https://github.com/dicarlolab/CORnet.

#### Estimation of discriminatory index (d’)

The discrimination index was computed to quantify the difference between the match of the ANNs’ (models per layer) behavioral predictions to the behavior measured in *Controls* and IwA (as shown in Figure 2E). It was calculated as:

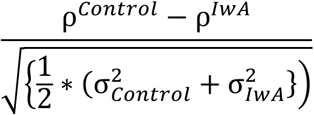

where ρ*^Control^* and ρ*^IwA^* was the correlation between ANN predictions and behavior measured in *Controls* and IwA respectively. σ*_Control_* and σ_IwA_ was the standard deviation of the bootstrap estimates of the correlations with random subsampling features from the model layers. To make the comparisons fair across all layers, 1000 features were randomly subsampled (without repetition) 100 times to estimate the ANN predictions.

#### Estimation of residuals between ANN-IT and human amygdala’s behavioral predictions

I first estimated the cross-validated test predictions (*ANN^Pred^*) of behavioral patterns from an ANN-IT layer (e.g., AlexNet ‘fc7’ model used in the study) using the partial least squares regression method. The ground truth values of image-level facial happiness were used as the dependent variable in this analysis. Next, I used the same algorithm but with the human amygdala neural features (instead of the ANN-IT features) as the predictors to estimate the neurally decoded behavioral patterns (*Amygdala^Pred^*). I then used a generalized linear regression model (MATLAB: *glmfit*) to estimate the residues while using *ANN^Pred^* as the predictor and *Amygdala^Pred^* as the dependent variable. The square of the Pearson correlation (%EV) between this residue vector (one value per image) and the image-level behavioral vector (Probability of choosing “Happy” per image) measured in the *Controls* is plotted in the y-axis of Figure 4 (left panels). These %EV values were corrected by the noise estimates in the behavioral data per image selection. In addition, all %EV values were estimated in a cross validated way, wherein the image selections and the final estimates were done based on different groups of subjects.

#### In silico model perturbation and training

*Generation of activity scaled additive noise values:* To estimate how much noise shall be added to each unit (feature) of the model layer, I used the following procedure. First, I estimated the standard deviation (σ, across all 28 images) of the activation distribution per unit in a noise-free model. The addition of noise was made proportional to this value. To vary noise levels, a scalar factor (*C*; x-axis in Figure5D and 5E) was multiplied with *σ* per unit. For each unit, the noise added was drawn from a normal distribution that had a standard deviation of *C*σ*.

*Training the model with and without noise:* To simulate a learning scheme with noise, I modified the model feature activations in the following way. During training of the regression model (i.e., estimating 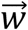 and *bias*), the noisy version of the model was generated by concatenating 1000 randomly drawn features (which were fixed for each iteration of the procedure), with ten repetitions of the same features but with the added noise on top of it. This procedure was repeated several times to estimate the variance in the model predictions per noise level. For the noise free model, the same 1000 randomly drawn features were repeated without addition of any noise.

### Statistics

All correlation scores reported in this study are Kendall rank coefficients (unless otherwise mentioned). For significance tests of correlations (between two variables of interest), I have used a bootstrapped permutation test. To do this, I first constructed a null hypothesis by mixing the two variables and then randomly drew (as many times as the number of elements in the original variable) with replacements two elements from the mixed dataset to create two vectors. These two vectors can be constructed multiple times (typically >100) and correlated. The resulting correlation distribution was considered as the null hypothesis. Then the true raw correlation was compared to this distribution to determine a p-value of rejecting the null distribution.

## Data and Code Availability

All the data and code used in this study will be freely available to download and use during the time of journal publication from https://github.com/kohitij-kar/2021_faceEmotion_ASD.

## Acknowledgments

I thank R. Adolphs, P. Sinha (and Sinha Lab members), and J.J. DiCarlo for helpful comments and discussions. I thank S. Wang for sharing the behavioral and neural datasets used in this study. I thank S. Wang, S. Sanghavi, A. Peter, and Y. Bai for helpful comments on the manuscript.

## Notes

### Competing Interest Statement

The authors have declared no competing interest.

